## Supplementary Material (pdf) for "Temporal origin of nestedness in interaction networks"

### *Supplementary Material*
### Temporal origin of nestedness in interaction networks

Phillip P. A. Staniczenko[1] and Debabrata Panja[2,3]

[1]Department of Biology, Brooklyn College, City University of New York, 2900 Bedford Avenue, Brooklyn, NY 11210, USA
[2]Department of Information and Computing Sciences, Utrecht University, Princetonplein 5, 3584 CC Utrecht, The Netherlands
[3]Centre for Complex Systems Studies, Utrecht University, The Netherlands

September 22, 2023

**Table S1.** Proportion of empirical nestedness generated by the degree distribution and phenology models at three levels of temporal aggregation: weekly, monthly, and annual.

**Table S2.** Phi coefficients and F-scores for the degree distribution and phenology models at three levels of temporal aggregation: weekly, monthly, and annual.

**Table S3.** Backbone of nestedness results for the phenology model at an annual level of temporal aggregation.

**Table S4.** Proportion of empirical nestedness, phi coefficients, and F-scores for the degree distribution, presence, and phenology models at an annual level of temporal aggregation.

**Figure S1.** Version of Figure 2 in the main paper with time series of nestedness values.

**Figure S2.** Histograms of the relative frequency of co-presences for auction market data.

**Figure S3.** Histograms of the relative frequency of co-presences for negotiation market data.

**Figure S4.** Histograms of the relative frequency of co-presences for plant-pollinator data.

**Table S1:** Proportion of empirical nestedness generated by the degree distribution ($\Delta\tilde{\rho}_{\mathrm{DD}}$) and phenology ($\Delta\tilde{\rho}_{\mathrm{Phen}}$) models at three levels of temporal aggregation: weekly (W), monthly (M), and annual (Y); mean and standard deviation (in parentheses) of 1000 realizations.

| Data set | Ref | Agg | $\Delta\tilde{\rho}_{\mathrm{DD}}$ | $\Delta\tilde{\rho}_{\mathrm{Phen}}$ |
|---|---|---|---|---|
| Auction | [1] | Y | 0.26 (0.02) | 0.91 (0.02) |
| | | M | 0.24 (0.03) | 0.53 (0.03) |
| | | W | 0.26 (0.05) | 0.30 (0.05) |
| Negotiation | [1] | Y | 0.26 (0.03) | 0.81 (0.03) |
| | | M | 0.27 (0.03) | 0.44 (0.03) |
| | | W | 0.28 (0.04) | 0.28 (0.04) |
| CaraDonna | [2] | Y | 0.29 (0.09) | 0.78 (0.10) |
| | | M | 0.32 (0.13) | 0.72 (0.14) |
| | | W | 0.43 (0.70) | 0.52 (0.74) |
| Alarcon2008 | [3] | Y | 0.36 (0.08) | 0.63 (0.18) |
| Benadi2014 | [4] | Y | 0.27 (0.02) | 0.83 (0.03) |
| | | M | 0.37 (0.03) | 0.52 (0.04) |
| Burkle2012 | [5] | Y | 0.28 (0.05) | 0.54 (0.05) |
| Chacoff2018 | [6] | Y | 0.31 (0.09) | 0.73 (0.10) |
| Fruend2010 | [7] | Y | 0.29 (0.05) | 1.03 (0.06) |
| | | M | 0.34 (0.13) | 0.92 (0.17) |
| LaraRomero2016 | [8] | Y | 0.30 (0.06) | 0.82 (0.07) |
| LeBuhnYY | [9] | Y | 0.29 (0.07) | 0.89 (0.08) |
| Olito2015 | [10] | Y | 0.33 (0.08) | 1.02 (0.09) |
| Rasmussen2013 | [11] | Y | 0.31 (0.09) | 1.01 (0.11) |
| ResascoYY | [9] | Y | 0.37 (0.10) | 0.71 (0.11) |
| Simanonok2014 | [12] | Y | 0.29 (0.06) | 0.93 (0.07) |
| Thompson2018 | [13] | Y | 0.35 (0.07) | 0.99 (0.08) |
| Vazquez2003 | [14] | Y | 0.44 (0.13) | 0.92 (0.12) |
| | | M | 0.67 (1.38) | 1.95 (3.71) |
| Weiner2014 | [15] | Y | 0.29 (0.02) | 0.81 (0.02) |
| | | M | 0.35 (0.05) | 0.62 (0.05) |
| | | W | 0.39 (0.10) | 0.42 (0.09) |
| Winfree2014 | [16] | Y | 0.31 (0.10) | 0.67 (0.11) |
| WinfreeYYa | [17] | Y | 0.30 (0.08) | 0.82 (0.09) |
| | | M | 0.48 (0.79) | 0.86 (1.21) |
| WinfreeYYb | [17] | Y | 0.29 (0.06) | 0.83 (0.07) |
| | | M | 0.43 (0.54) | 0.76 (0.61) |
| WinfreeYYc | [9] | Y | 0.35 (0.14) | 0.50 (0.14) |
| WinfreeYYd | [18] | Y | 0.26 (0.03) | 0.80 (0.03) |
| | | M | 0.27 (0.05) | 0.63 (0.05) |
| | | W | 0.31 (0.11) | 0.40 (0.11) |
| WinfreeYYe | [9] | Y | 0.30 (0.07) | 0.96 (0.10) |
| | | M | 0.30 (0.08) | 0.96 (0.10) |
| | | W | 0.41 (0.39) | 0.75 (0.43) |
| WinfreeYYf | [19] | Y | 0.27 (0.06) | 0.54 (0.07) |
| | | M | 0.38 (0.29) | 0.51 (0.29) |

**Table S2:** Phi coefficients ($\phi$) and F-scores ($F_1$) for the degree distribution (DD) and phenology (Phen) models at three levels of temporal aggregation: weekly (W), monthly (M), and annual (Y); mean and standard deviation (in parentheses) of 1,000 realizations.

| Data set | Ref | Agg | $\phi_{DD}$ | $\phi_{Phen}$ | $F_{1DD}$ | $F_{1Phen}$ |
|---|---|---|---|---|---|---|
| Auction | [1] | Y | 0.25 (0.01) | 0.47 (0.01) | 0.53 (0.01) | 0.67 (0.00) |
| | | M | 0.20 (0.01) | 0.29 (0.01) | 0.41 (0.01) | 0.47 (0.01) |
| | | W | 0.17 (0.02) | 0.14 (0.02) | 0.35 (0.01) | 0.33 (0.01) |
| Negotiation | [1] | Y | 0.23 (0.01) | 0.35 (0.01) | 0.61 (0.00) | 0.68 (0.00) |
| | | M | 0.17 (0.01) | 0.16 (0.01) | 0.38 (0.01) | 0.37 (0.01) |
| | | W | 0.13 (0.01) | 0.08 (0.01) | 0.26 (0.01) | 0.22 (0.01) |
| CaraDonna | [2] | Y | 0.14 (0.03) | 0.34 (0.03) | 0.26 (0.02) | 0.44 (0.02) |
| | | M | 0.17 (0.04) | 0.28 (0.04) | 0.31 (0.03) | 0.41 (0.03) |
| | | W | 0.20 (0.08) | 0.16 (0.07) | 0.38 (0.06) | 0.35 (0.06) |
| Alarcon2008 | [3] | Y | 0.10 (0.02) | 0.17 (0.02) | 0.16 (0.02) | 0.22 (0.02) |
| Benadi2014 | [4] | Y | 0.07 (0.01) | 0.19 (0.01) | 0.10 (0.01) | 0.22 (0.01) |
| | | M | 0.08 (0.02) | 0.14 (0.02) | 0.14 (0.02) | 0.19 (0.02) |
| Burkle2012 | [5] | Y | 0.12 (0.02) | 0.19 (0.02) | 0.19 (0.02) | 0.25 (0.02) |
| Chacoff2018 | [6] | Y | 0.11 (0.02) | 0.19 (0.03) | 0.19 (0.02) | 0.27 (0.02) |
| Fruend2010 | [7] | Y | 0.09 (0.01) | 0.28 (0.02) | 0.14 (0.01) | 0.32 (0.02) |
| | | M | 0.08 (0.05) | 0.21 (0.05) | 0.16 (0.04) | 0.28 (0.05) |
| LaraRomero2016 | [8] | Y | 0.15 (0.02) | 0.31 (0.02) | 0.29 (0.02) | 0.42 (0.02) |
| LeBuhnYY | [9] | Y | 0.07 (0.02) | 0.29 (0.02) | 0.12 (0.02) | 0.33 (0.02) |
| Olito2015 | [10] | Y | 0.09 (0.02) | 0.26 (0.02) | 0.14 (0.02) | 0.30 (0.02) |
| Rasmussen2013 | [11] | Y | 0.10 (0.02) | 0.28 (0.02) | 0.19 (0.02) | 0.35 (0.02) |
| ResascoYY | [9] | Y | 0.10 (0.02) | 0.20 (0.03) | 0.17 (0.02) | 0.26 (0.02) |
| Simanonok2014 | [12] | Y | 0.10 (0.02) | 0.26 (0.02) | 0.16 (0.02) | 0.31 (0.02) |
| Thompson2018 | [13] | Y | 0.11 (0.02) | 0.26 (0.02) | 0.19 (0.02) | 0.33 (0.02) |
| Vazquez2003 | [14] | Y | 0.13 (0.03) | 0.49 (0.03) | 0.24 (0.03) | 0.55 (0.02) |
| | | M | 0.24 (0.10) | 0.39 (0.09) | 0.45 (0.06) | 0.55 (0.06) |
| Weiner2014 | [15] | Y | 0.11 (0.01) | 0.28 (0.01) | 0.15 (0.01) | 0.31 (0.01) |
| | | M | 0.12 (0.02) | 0.20 (0.01) | 0.17 (0.01) | 0.25 (0.02) |
| | | W | 0.13 (0.03) | 0.14 (0.03) | 0.21 (0.02) | 0.21 (0.02) |
| Winfree2014 | [16] | Y | 0.09 (0.02) | 0.17 (0.03) | 0.17 (0.02) | 0.24 (0.02) |
| WinfreeYYa | [17] | Y | 0.11 (0.02) | 0.28 (0.03) | 0.18 (0.02) | 0.34 (0.02) |
| | | M | 0.12 (0.05) | 0.21 (0.05) | 0.23 (0.04) | 0.31 (0.04) |
| WinfreeYYb | [17] | Y | 0.09 (0.02) | 0.25 (0.02) | 0.15 (0.02) | 0.30 (0.02) |
| | | M | 0.10 (0.05) | 0.21 (0.05) | 0.21 (0.04) | 0.30 (0.04) |
| WinfreeYYc | [9] | Y | 0.23 (0.05) | 0.24 (0.05) | 0.39 (0.04) | 0.40 (0.04) |
| WinfreeYYd | [18] | Y | 0.16 (0.01) | 0.35 (0.01) | 0.26 (0.01) | 0.42 (0.01) |
| | | M | 0.16 (0.02) | 0.25 (0.02) | 0.26 (0.02) | 0.35 (0.02) |
| | | W | 0.14 (0.03) | 0.15 (0.03) | 0.25 (0.03) | 0.26 (0.03) |
| WinfreeYYe | [9] | Y | 0.12 (0.03) | 0.27 (0.03) | 0.23 (0.02) | 0.36 (0.02) |
| | | M | 0.12 (0.03) | 0.28 (0.03) | 0.23 (0.02) | 0.36 (0.02) |
| | | W | 0.11 (0.04) | 0.15 (0.05) | 0.22 (0.04) | 0.26 (0.04) |
| WinfreeYYf | [19] | Y | 0.08 (0.02) | 0.23 (0.02) | 0.15 (0.02) | 0.29 (0.02) |
| | | M | 0.11 (0.05) | 0.19 (0.05) | 0.23 (0.04) | 0.29 (0.04) |

**Table S3:** Backbone of nestedness results for the phenology model at an annual level of temporal aggregation: (i) ratio of nestedness values for the subsets of true positive and false negative links, $\rho_{\mathrm{TP}}/\rho_{\mathrm{FN}}$ and (ii) ratio of the deviations of nestedness relative to comparable Erdös-Rényi random graphs, $\rho_{\mathrm{TP},\Delta\mathrm{ER}}/\rho_{\mathrm{FN},\mathrm{ER}}$, where $\rho_{\mathrm{TP},\Delta\mathrm{ER}} = (\rho_{\mathrm{TP}} - \rho_{\mathrm{TP},\mathrm{ER}})/(1 - \rho_{\mathrm{TP},\mathrm{ER}})$ is the proportional deviation for the subset of true positive links and $\rho_{\mathrm{FN},\mathrm{ER}} = (\rho_{\mathrm{FN}} - \rho_{\mathrm{FN},\mathrm{ER}})/(1 - \rho_{\mathrm{FN},\ \mathrm{RG}})$ is the proportional deviation for the subset of false negative links; mean and standard deviation (in parentheses) of 1000 realizations.

| Data set | Ref | $\rho_{\mathrm{TP}}/\rho_{\mathrm{FN}}$ | $\rho_{\mathrm{TP},\Delta\mathrm{RG}}/\rho_{\mathrm{FN},\Delta\mathrm{RG}}$ |
|---|---|---|---|
| Auction | [1] | 1.64 (0.02) | 2.60 (0.17) |
| Negotiation | [1] | 1.55 (0.01) | 2.56 (0.23) |
| CaraDonna | [2] | 1.29 (0.06) | 2.02 (0.64) |
| Alarcon2008 | [3] | 1.26 (0.09) | 0.92 (0.48) |
| Benadi2014 | [4] | 1.27 (0.04) | 1.24 (0.12) |
| Burkle2012 | [5] | 1.16 (0.06) | 0.89 (0.23) |
| Chacoff2018 | [6] | 1.17 (0.07) | 0.77 (0.48) |
| Fruend2010 | [7] | 1.38 (0.06) | 1.56 (0.25) |
| LaraRomero2016 | [8] | 1.31 (0.04) | 2.25 (0.50) |
| LeBuhnYY | [9] | 1.46 (0.08) | 2.04 (0.54) |
| Olito2015 | [10] | 1.46 (0.07) | 1.98 (0.49) |
| Rasmussen2013 | [11] | 1.37 (0.07) | 2.25 (0.83) |
| ResascoYY | [9] | 1.22 (0.08) | 0.86 (0.56) |
| Simanonok2014 | [12] | 1.38 (0.07) | 1.77 (0.40) |
| Thompson2018 | [13] | 1.22 (0.06) | 1.55 (0.38) |
| Vazquez2003 | [14] | 1.07 (0.06) | 1.60 (0.76) |
| Weiner2014 | [15] | 1.29 (0.03) | 1.31 (0.08) |
| Winfree2014 | [16] | 1.32 (0.10) | 1.58 (0.60) |
| WinfreeYYa | [17] | 1.28 (0.07) | 1.46 (0.44) |
| WinfreeYYb | [17] | 1.18 (0.06) | 0.96 (0.27) |
| WinfreeYYc | [9] | 1.13 (0.07) | 0.97 (0.85) |
| WinfreeYYd | [18] | 1.41 (0.04) | 1.77 (0.17) |
| WinfreeYYe | [9] | 1.33 (0.06) | 1.73 (0.51) |
| WinfreeYYf | [19] | 1.09 (0.06) | 0.75 (0.28) |

**Table S4:** Proportion of empirical nestedness ($\Delta\tilde{\rho}$), phi coefficients ($\phi$), and F-scores ($F_1$) for the degree distribution (DD), presence (Pres), and phenology (Phen) models at an annual level of temporal aggregation, mean and standard deviation (in parentheses) of 1000 realizations.

| Data set | Ref | $\Delta\tilde{\rho}_{DD}$ | $\Delta\tilde{\rho}_{Pres}$ | $\Delta\tilde{\rho}_{Phen}$ | $\phi_{DD}$ | $\phi_{Pres}$ | $\phi_{Phen}$ | $F_{1DD}$ | $F_{1Pres}$ | $F_{1Phen}$ |
|---|---|---|---|---|---|---|---|---|---|---|
| Auction | [1] | 0.26 (0.02) | 1.04 (0.02) | 0.91 (0.02) | 0.25 (0.01) | 0.31 (0.01) | 0.47 (0.01) | 0.53 (0.01) | 0.58 (0.00) | 0.67 (0.00) |
| Negotiation | [1] | 0.26 (0.03) | 0.37 (0.03) | 0.81 (0.03) | 0.23 (0.01) | 0.22 (0.01) | 0.35 (0.01) | 0.61 (0.00) | 0.61 (0.00) | 0.68 (0.00) |
| CaraDonna | [2] | 0.30 (0.09) | 0.59 (0.09) | 0.78 (0.10) | 0.14 (0.03) | 0.25 (0.03) | 0.34 (0.03) | 0.26 (0.02) | 0.36 (0.02) | 0.44 (0.02) |
| Alarcon2008 | [3] | 0.36 (0.08) | 0.56 (0.08) | 0.63 (0.08) | 0.10 (0.02) | 0.14 (0.02) | 0.17 (0.02) | 0.16 (0.02) | 0.20 (0.02) | 0.22 (0.02) |
| Benadi2014 | [4] | 0.27 (0.02) | 0.59 (0.03) | 0.83 (0.03) | 0.07 (0.01) | 0.14 (0.01) | 0.19 (0.01) | 0.10 (0.01) | 0.17 (0.01) | 0.22 (0.01) |
| Burkle2012 | [5] | 0.28 (0.05) | 0.47 (0.06) | 0.54 (0.05) | 0.12 (0.02) | 0.15 (0.02) | 0.19 (0.02) | 0.19 (0.02) | 0.22 (0.02) | 0.25 (0.02) |
| Chacoff2018 | [6] | 0.31 (0.09) | 0.58 (0.09) | 0.73 (0.10) | 0.11 (0.02) | 0.14 (0.02) | 0.19 (0.03) | 0.19 (0.02) | 0.22 (0.02) | 0.27 (0.02) |
| Fruend2010 | [7] | 0.29 (0.05) | 0.84 (0.06) | 1.03 (0.06) | 0.09 (0.01) | 0.24 (0.02) | 0.28 (0.02) | 0.14 (0.01) | 0.28 (0.02) | 0.32 (0.02) |
| LaraRomero2016 | [8] | 0.30 (0.06) | 0.51 (0.06) | 0.82 (0.07) | 0.15 (0.02) | 0.23 (0.02) | 0.31 (0.02) | 0.29 (0.02) | 0.35 (0.02) | 0.42 (0.02) |
| LeBuhnYY | [9] | 0.29 (0.07) | 1.01 (0.08) | 0.89 (0.08) | 0.07 (0.02) | 0.25 (0.02) | 0.29 (0.02) | 0.12 (0.02) | 0.29 (0.02) | 0.33 (0.02) |
| Olito2015 | [10] | 0.33 (0.08) | 0.80 (0.08) | 1.02 (0.09) | 0.09 (0.02) | 0.20 (0.02) | 0.26 (0.02) | 0.14 (0.02) | 0.25 (0.02) | 0.30 (0.02) |
| Rasmussen2013 | [11] | 0.31 (0.09) | 0.83 (0.10) | 1.01 (0.11) | 0.10 (0.02) | 0.21. (0.02) | 0.28 (0.02) | 0.19 (0.02) | 0.29 (0.02) | 0.35 (0.02) |
| ResascoYY | [9] | 0.37 (0.10) | 0.66 (0.11) | 0.71 (0.11) | 0.10 (0.02) | 0.17 (0.03) | 0.20 (0.03) | 0.17 (0.02) | 0.23 (0.02) | 0.26 (0.02) |
| Simanonok2014 | [12] | 0.29 (0.06) | 0.72 (0.07) | 0.93 (0.07) | 0.10 (0.02) | 0.21 (0.02) | 0.26 (0.02) | 0.16 (0.02) | 0.27 (0.02) | 0.31 (0.02) |
| Thompson2018 | [13] | 0.35 (0.07) | 1.02 (0.08) | 0.99 (0.08) | 0.11 (0.02) | 0.24 (0.02) | 0.26 (0.02) | 0.19 (0.02) | 0.31 (0.02) | 0.33 (0.02) |
| Vazquez2003 | [14] | 0.44 (0.13) | 0.85 (0.11) | 0.92 (0.12) | 0.13 (0.03) | 0.46 (0.03) | 0.49 (0.03) | 0.24 (0.03) | 0.53 (0.03) | 0.55 (0.02) |
| Weiner2014 | [15] | 0.29 (0.02) | 0.58 (0.02) | 0.81 (0.02) | 0.11 (0.01) | 0.21 (0.01) | 0.28 (0.01) | 0.15 (0.01) | 0.24 (0.01) | 0.31 (0.01) |
| Winfree2014 | [16] | 0.31 (0.10) | 0.67 (0.11) | 0.67 (0.11) | 0.09 (0.02) | 0.16 (0.03) | 0.17 (0.03) | 0.17 (0.02) | 0.24 (0.02) | 0.24 (0.02) |
| WinfreeYYa | [17] | 0.30 (0.08) | 0.87 (0.09) | 0.82 (0.09) | 0.11 (0.02) | 0.26 (0.03) | 0.28 (0.03) | 0.18 (0.02) | 0.32 (0.02) | 0.34 (0.02) |
| WinfreeYYb | [17] | 0.29 (0.06) | 0.89 (0.07) | 0.83 (0.07) | 0.09 (0.02) | 0.22 (0.02) | 0.25 (0.02) | 0.15 (0.02) | 0.28 (0.02) | 0.30 (0.02) |
| WinfreeYYc | [9] | 0.35 (0.14) | 0.33 (0.13) | 0.50 (0.14) | 0.23 (0.05) | 0.16 (0.05) | 0.24 (0.05) | 0.39 (0.04) | 0.33 (0.04) | 0.40 (0.04) |
| WinfreeYYd | [18] | 0.26 (0.03) | 0.50 (0.03) | 0.80 (0.03) | 0.16 (0.01) | 0.22 (0.01) | 0.35 (0.01) | 0.26 (0.01) | 0.31 (0.01) | 0.42 (0.01) |
| WinfreeYYe | [9] | 0.30 (0.07) | 0.64 (0.09) | 0.96 (0.10) | 0.12 (0.03) | 0.19 (0.03) | 0.27 (0.03) | 0.23 (0.02) | 0.29 (0.02) | 0.36 (0.02) |
| WinfreeYYf | [19] | 0.27 (0.06) | 0.84 (0.09) | 0.54 (0.07) | 0.08 (0.02) | 0.23 (0.02) | 0.23 (0.02) | 0.15 (0.02) | 0.29 (0.02) | 0.29 (0.02) |

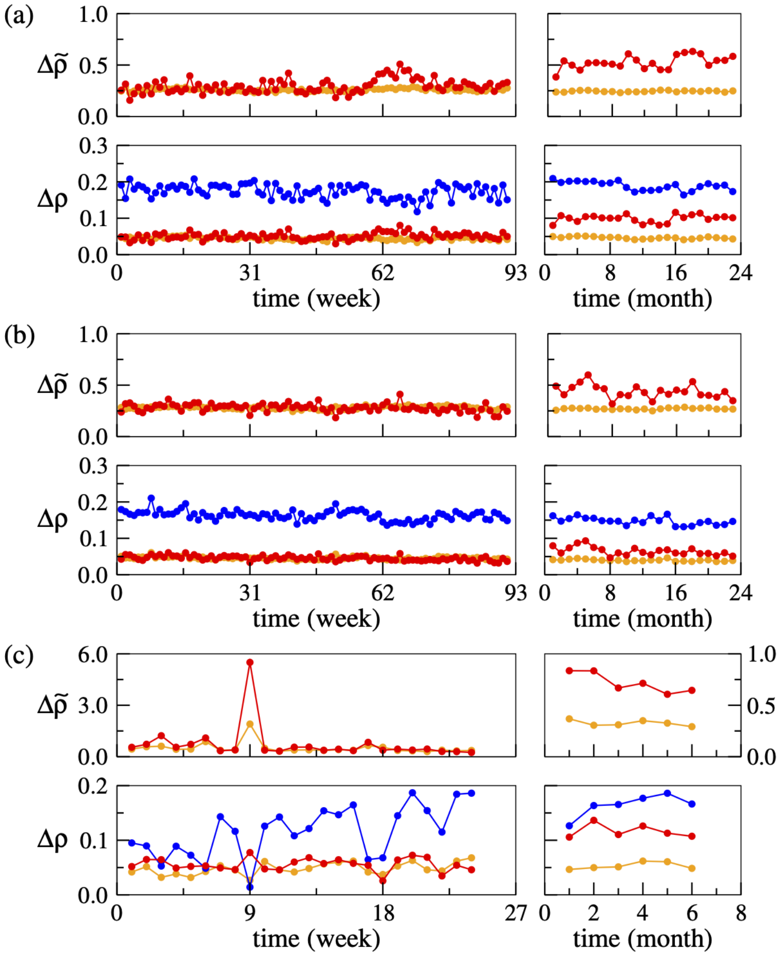


| year | 1 | 2 |
|---|---|---|
| $\Delta\tilde{\rho}$ | 0.923<br>0.259 | 0.893<br>0.266 |
| $\Delta\rho$ | 0.160<br>0.148<br>0.041 | 0.145<br>0.130<br>0.038 |

| year | 1 | 2 |
|---|---|---|
| $\Delta\tilde{\rho}$ | 0.773<br>0.262 | 0.851<br>0.257 |
| $\Delta\rho$ | 0.101<br>0.078<br>0.027 | 0.108<br>0.092<br>0.028 |

| year | 1 | 2 | 3 |
|---|---|---|---|
| $\Delta\tilde{\rho}$ | 0.837<br>0.313 | 0.828<br>0.288 | 0.685<br>0.289 |
| $\Delta\rho$ | 0.140<br>0.117<br>0.044 | 0.201<br>0.167<br>0.058 | 0.181<br>0.124<br>0.052 |

**Figure S1:** Nestedness generated by the degree distribution and phenology models relative to empirical networks built from three data sets — buyer-seller transactions in auction (a) and negotiation (b) fish markets in Boulogne-sur-Mer, France, and plant-pollinator visitations in field sites at Rocky Mountain Biological Laboratory, Colorado, USA (c) — at three levels of temporal aggregation: week (left), month (middle), and year (right). In the lower plots of each panel, nestedness values for empirical networks (blue), degree distribution model networks (orange), and phenology model networks (red) are measured relative to corresponding values for Erdös-Rényi random graphs ($\Delta\rho = 0$); in the upper plots, nestedness values for the two models are scaled between corresponding values for Erdös-Rényi random graphs ($\Delta\tilde{\rho} = 0$) and empirical networks ($\Delta\tilde{\rho} = 1$). While the levels of nestedness generated by the degree distribution model remain relatively low as temporal aggregation increases, the phenology model generates values that approach empirically observed levels.

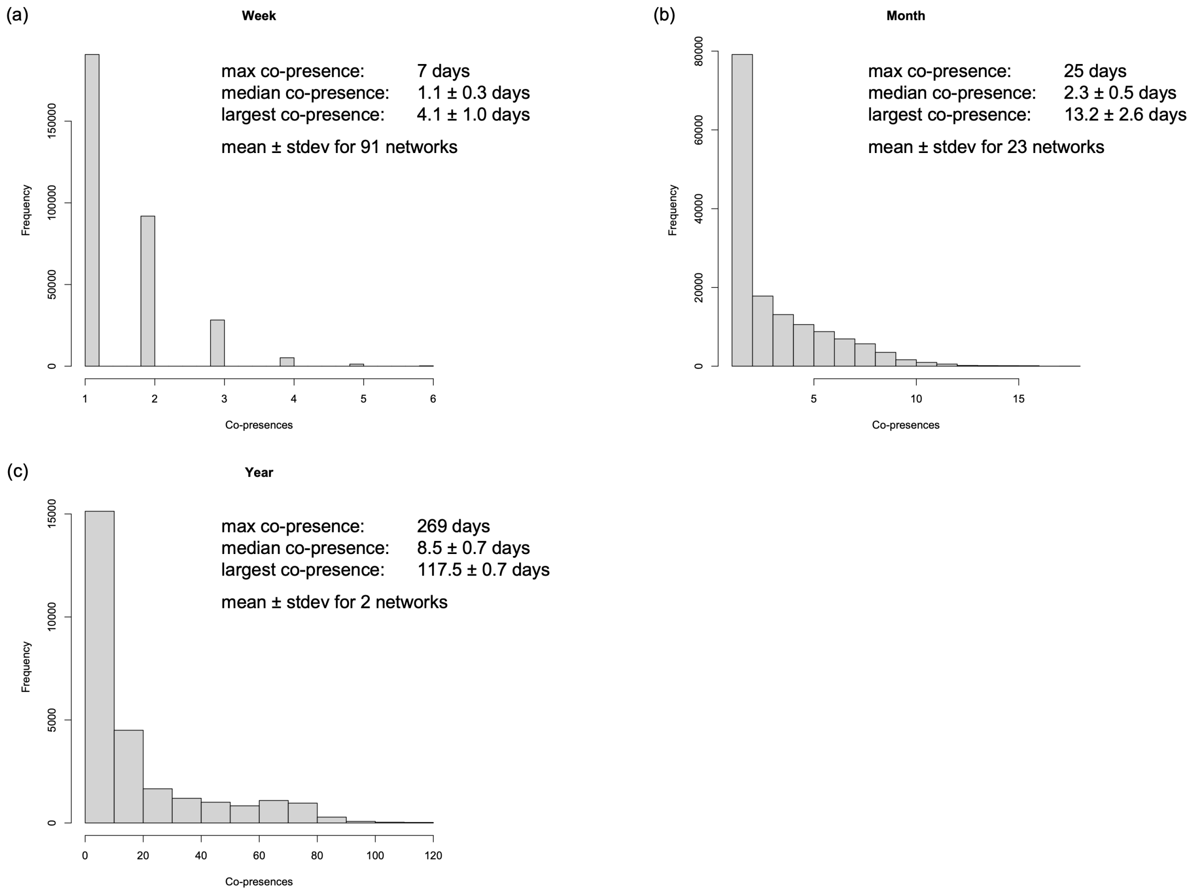


**Figure S2:** Histograms of the relative frequency of co-presences for auction market data [1]. Panels show co-presences for all networks combined at the specific level of temporal aggregation while statistics for the median and largest number of co-presences statistics present averages and standard deviations for individual networks.

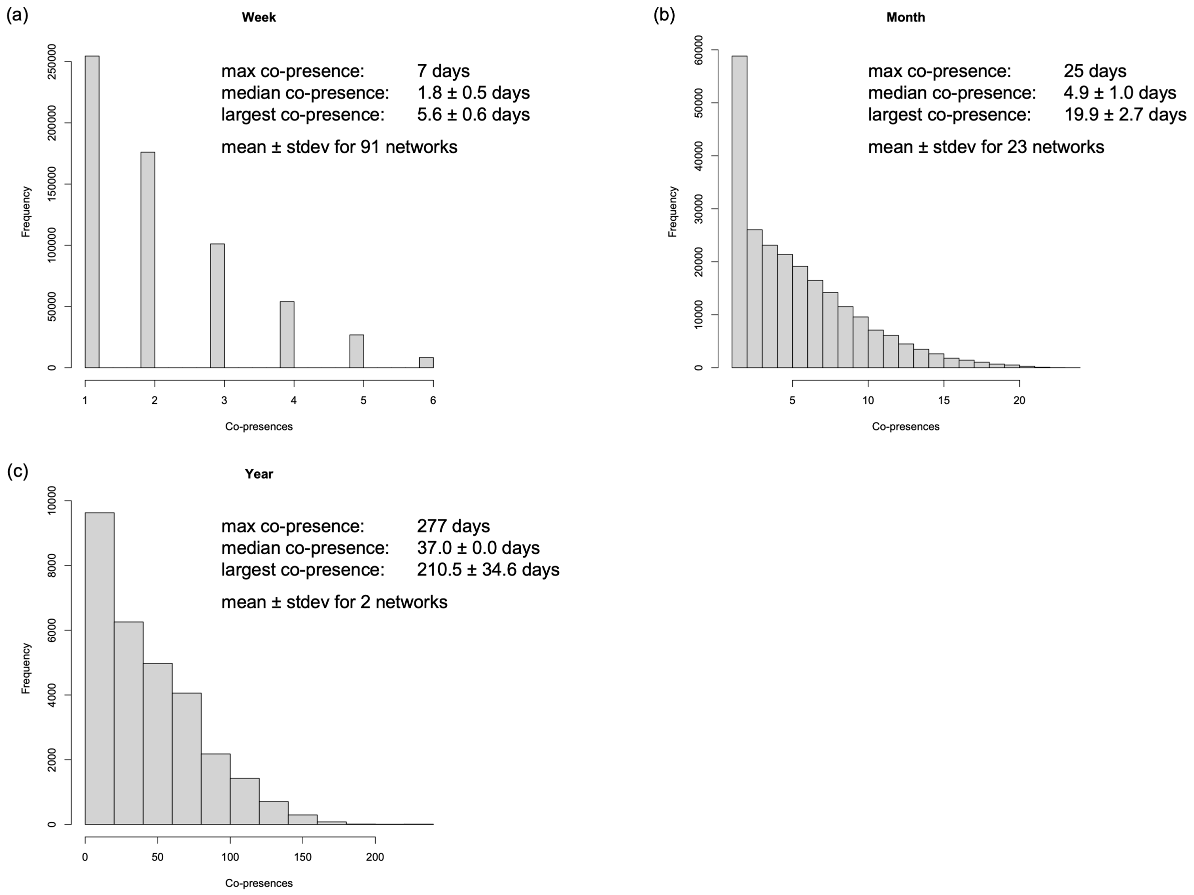


**Figure S3:** Histograms of the relative frequency of co-presences for negotiation market data [1]. Panels show co-presences for all networks combined at the specific level of temporal aggregation while statistics for the median and largest number of co-presences statistics present averages and standard deviations for individual networks.

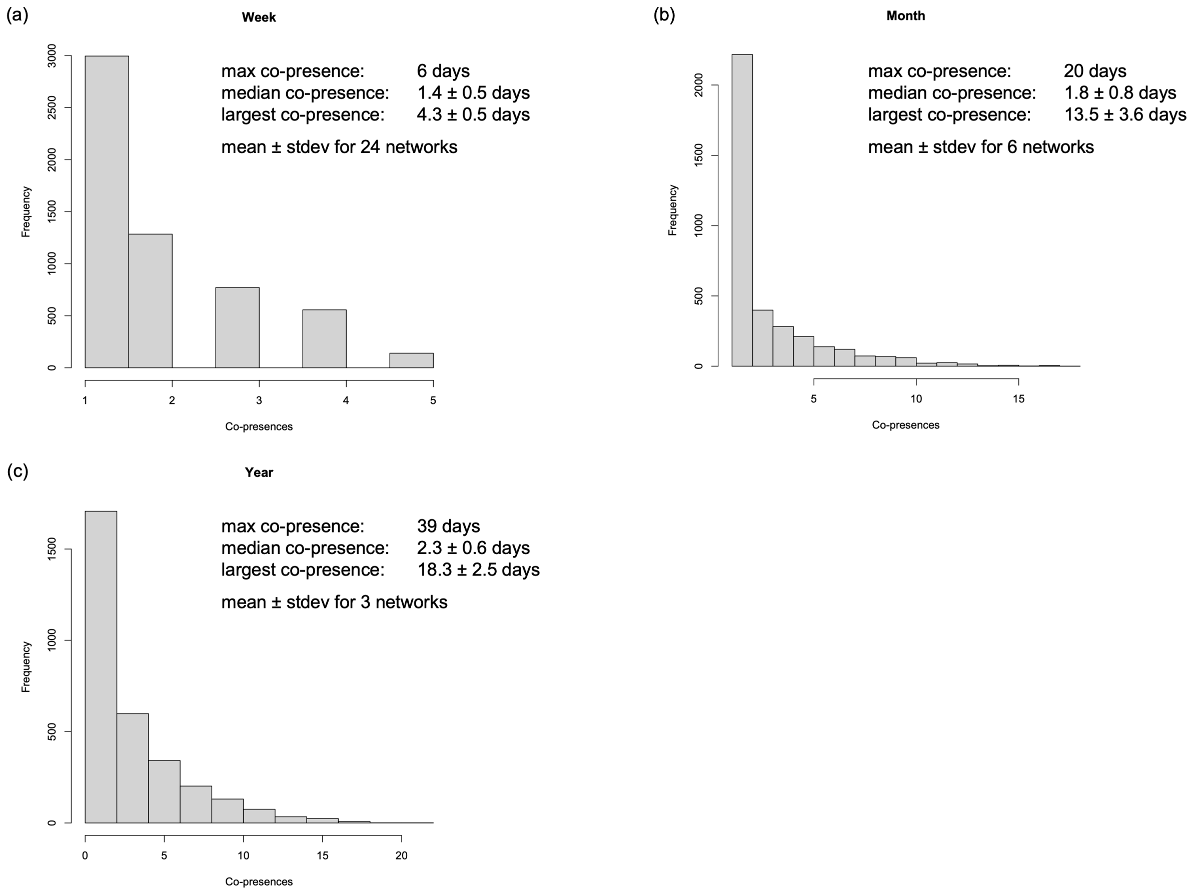


**Figure S4:** Histograms of the relative frequency of co-presences for plant-pollinator data [2]. Panels show co-presences for all networks combined at the specific level of temporal aggregation while statistics for the median and largest number of co-presences statistics present averages and standard deviations for individual networks.
